# An evolutionary conserved division-of-labor between hippocampal and neocortical sharp-wave ripples organizes information transfer during sleep

**DOI:** 10.1101/2022.10.19.512822

**Authors:** Frank J. van Schalkwijk, Jan Weber, Michael A. Hahn, Janna D. Lendner, Marion Inostroza, Jack J. Lin, Randolph F. Helfrich

## Abstract

The hippocampal sharp-wave ripple (SW-R) is the key substrate of the hippocampal-neocortical dialogue underlying memory formation. Recently, it became evident that SW-R are not unique to archicortex, but constitute a wide-spread neocortical phenomenon. To date, little is known about morphological and functional similarities between archi- and neocortical SW-R. Leveraging intracranial recordings from the human hippocampus and prefrontal cortex during sleep, our results reveal region-specific functional specializations, albeit a near-uniform morphology. While hippocampal SW-R trigger directional hippocampal-to-neocortical information flow, neocortical SW-R reduce information flow to minimize interference. At the population level, hippocampal SW-R confined population dynamics to a low-dimensional subspace, while neocortical SW-R diversified the population response; functionally uncoupling the hippocampal-neocortical network. Critically, our replication in rodents demonstrated the same division-of-labor between archi-and neocortical SW-R. These results uncover an evolutionary preserved mechanism where coordinated interplay between hippocampal and neocortical SW-R temporally segregates hippocampal information transfer from neocortical processing.

## Introduction

Sharp-wave ripple oscillations (SW-R; ~100-200 Hz) hallmark the hippocampal-neocortical dialogue that underlies sleep-dependent memory consolidation (Buzsaki, 2015). The systems memory consolidation theory suggests that SW-R synchronize large-scale population activity to reactivate mnemonic representations in the hippocampus, which are subsequently transferred to the neocortex for permanent storage (Frankland & Bontempi, 2005). This influential model further posits that SW-R do not occur in isolation, but are nested in other cardinal sleep oscillations, including cortical slow oscillations (< 1.25 Hz; SO) and thalamocortical spindles (~12-16 Hz; Buzsaki, 2015; Diekelmann & Born, 2010); thus, forming an oscillatory hierarchy that constitutes an endogenous timing mechanism to coordinate the information transfer from short- to long-term mnemonic storage. In support of key model assumptions, there is now mounting evidence that demonstrates a causal role of the hippocampal SW-R for memory formation (Ego-Stengel & Wilson, 2010; Girardeau et al., 2009; Maingret et al., 2016).

Importantly, the systems memory consolidation theory considers the hippocampus to be the sole source of SW-R. In contrast to this established notion, several recent studies demonstrated that SW-R are also prevalent in the neocortex. For example, parietal SW-R synchronize with their hippocampal counterparts during post-learning sleep in rodents (Khodagholy et al., 2017). Moreover, SW-R occur in most neocortical regions in humans (Dickey, Verzhbinsky, Jiang, Rosen, Kajfez, Eskandar, et al., 2022), with co-occurrence of hippocampal and neocortical SW-R being predictive of memory formation (Dickey, Verzhbinsky, Jiang, Rosen, Kajfez, Stedelin, et al., 2022; Vaz et al., 2019). Critically, these studies conceptualized hippocampal and neocortical SW-R as a unitary phenomenon given their striking morphological resemblance. To date, it is not known why such highly stereotypical electrophysiological signatures as SW-R emerge from the hippocampus and neocortex, despite clear differences in the underlying neuroanatomy. Especially the human PFC is often conceptualized as a key structure to house long-term mnemonic representations (Frankland & Bontempi, 2005); a structure where the rodent homologue is actively being debated (Carlen, 2017; Laubach et al., 2018). As a consequence, it remains unclear if neocortical SW-R fulfill the very same functional role as hippocampal SW-R in the human and rodent brain. Moreover, it is not known how SW-R that emerge from distinct anatomical structures interact to jointly coordinate the timed information transfer from short-to long-term storage.

Currently, the majority of evidence supporting the systems memory consolidation hypothesis stems from rodents, given the difficulty to non-invasively image the human hippocampus with a sufficient high spatiotemporal resolution to detect SW-R. Recently, several studies employing intracranial recordings in epileptic patients who undergo presurgical evaluation with implanted depth electrodes revealed that key principles of hippocampal-neocortical dialogue as observed in rodents, such as SO-spindle-ripple coupling, also apply to the human brain (Clemens et al., 2007; Helfrich et al., 2019; Skelin et al., 2021; Staresina et al., 2015). A major concern for human studies is that several electrophysiological phenomena that are inherent to the epileptic brain, such as interictal discharges (IED) or high-frequency oscillations (HFO) might confound SW-R detection (Frauscher et al., 2018; Jiang et al., 2020). While different algorithmic solutions have been introduced in the past to mitigate the effects of the underlying brain pathology (Norman et al., 2019; Skelin et al., 2021; Vaz et al., 2019), to date no study has validated their approach in rodents.

In the present study, we addressed these unanswered questions by directly recording from the hippocampus and adjacent structures in the medial temporal lobe (MTL) as well as the prefrontal cortex (PFC) in humans and rodents. We initially tested if SW-R are an inherent signature of the human PFC during sleep. Next, we determined whether SW-R in different network nodes fulfill distinct functional roles. Specifically, we assessed how hippocampal and neocortical SW-R modulate and synchronize population activity to support directed information transfer from putative short-term storage in the hippocampus to long-term storage in the neocortex. Lastly, employing a comparative approach, we determined whether findings in humans generalize to the rodent brain and whether the observed PFC signatures reflect true homologues of the hippocampal SW-R.

## Results

We combined intracranial recordings from the human MTL (*N_electrodes_* = 82) and PFC (*N_electrodes_* = 339) during sleep with polysomnography in patients with pharmacoresistant epilepsy during pre-surgical evaluation (*N* = 14; *M_age_* = 36.79 ± 3.55 years, mean ± SEM, range 19 to 58 years; 9 female; see **Fig. 1a** and ***SI Appendix, Table S*1** and **S2** for sleep architecture, spectral dynamics, and subject descriptives) to assess SW-R dynamics in support of network organization and information transfer during NREM sleep. Candidate SW-R were identified per electrode (Fig. 1B; Methods; Helfrich et al., 2019; Norman et al., 2019; Vaz et al., 2019). The ripple peak frequency was determined per region analogous to previous work in rodents (Bragin et al., 1999; Buzsaki, 2015) by considering a wide frequency range for event detection (60 – 240 Hz; ***SI Appendix*, Fig. S1**). Peak frequency for MTL (90.2 ± .03 Hz, mean ± SEM) and PFC ripples (93.3 ± .02 Hz) overlapped with the previously described frequency range (80-120 Hz) in humans (Bragin et al., 1999; Helfrich et al., 2019; Norman et al., 2021; Norman et al., 2019; Staresina et al., 2015) and hence, further analyses were subsequently constrained to this range (**Fig. 1b**). Artifactual events were rigorously excluded based on a stringent set of selection criteria (**Fig. 1c**; ***SI Appendix*, SI Text and Fig. S2**): (i) peak frequency in the human ripple range (80-120 Hz; Axmacher et al., 2008; Norman et al., 2019; Skelin et al., 2021; Staresina et al., 2015; Vaz et al., 2019); (ii) oscillatory characteristics of the ripple in the *time-domain:* multiple, evenly spaced distinct peaks; (iii) oscillatory characteristic of the ripple in the *frequency-domain:* transient, frequency-specific power increase (offset from the aperiodic 1/f component in the ripple frequency range); (iv) ripple nesting in the trough of a sharp-wave and (v) exclusion of sharp transients and inter-ictal discharges (IEDs). Application of these selection criteria yielded candidate SW-R events that exhibited a stereotypical morphology in the time-domain, both on the single-subject (**Fig. 1d and e**) as well as on the group level (see **Fig. 1f** for electrode coverage) for the MTL (**Fig. 1g**) and PFC (**Fig. 1h**). As indicated in the time-domain, the oscillatory characteristics of ripples were confirmed in the frequency-domain as isolated peaks in the power spectrum that rise above the 1/*f*^x^ background activity (**Fig. 1d and e,** *Center;* **Fig. 1g and h,** *Top Right*). The presence of a distinct oscillatory peak in the ripple band delineates SW-R from pathologic activity that results in a broadband power increase (***SI Appendix,*** **Fig. S2b and c**). Moreover, ripples were nested in the trough of the accompanying sharp wave component of SW-R (insets **Fig. 1g and h**) as demonstrated by significant coupling in the MTL (188.4° ± 6.76°; mean ± SEM; *p* < .0001, *Z* = 11.40, Rayleigh test; **Fig. 1g**, *Inset*) and PFC (165.0° ± 2.7°; *p* < .0001, Z = 13.56; **Fig. 1h**, *Inset*). Lastly, the power increase in the ripple frequency band was transient and confined to the duration of the sharp-wave ripple (**Fig. 1d and e,** *Right;* **Fig. 1g and h,** *Bottom Right*). Collectively, these findings establish that the detected events constitute the human homologues of SW-R that have previously been observed in the rodent hippocampus.

**Fig. 1.**
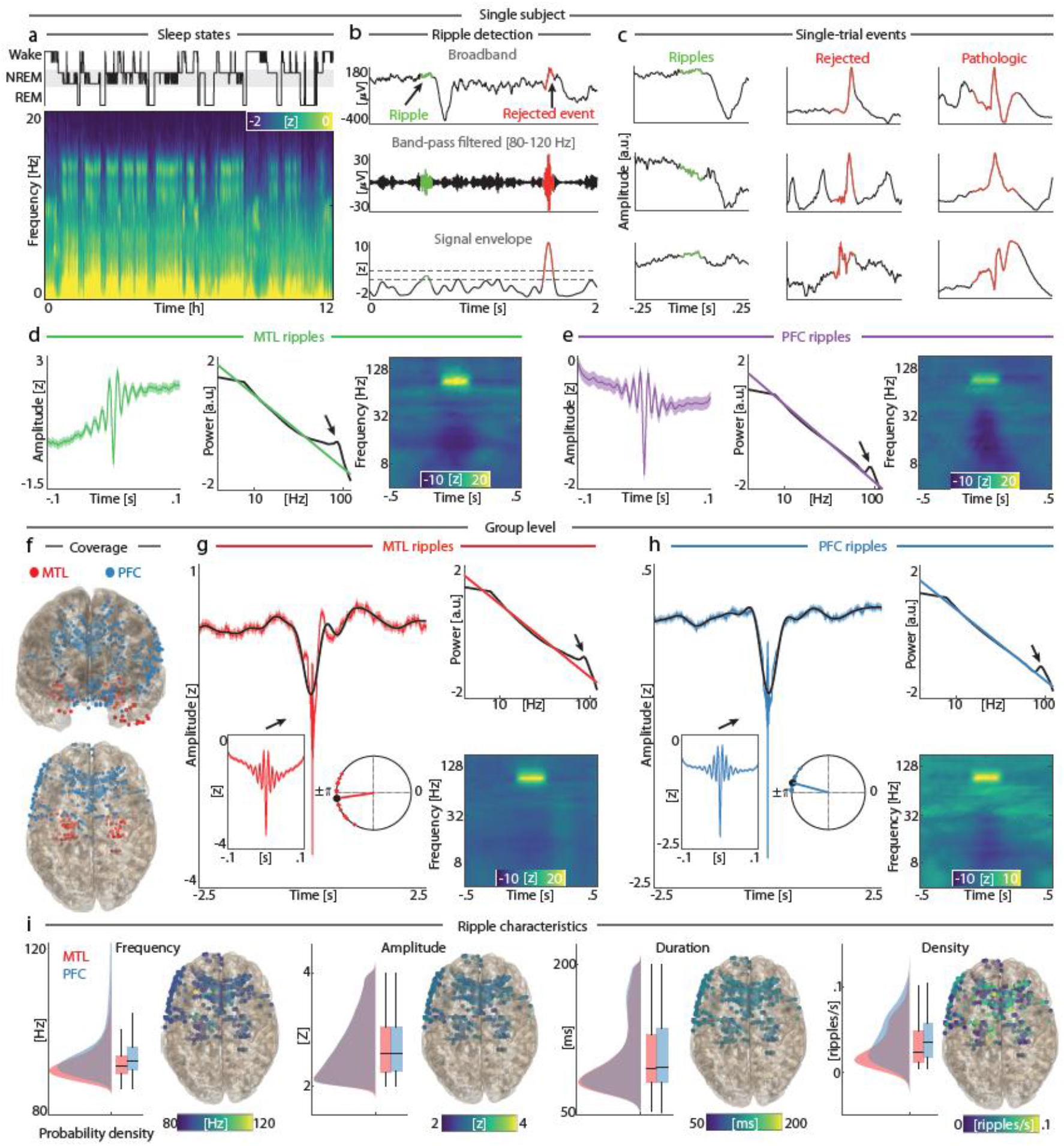
Ripple morphology in the MTL and PFC. (**a**) Sleep architecture of a single subject during a nighttime recording. *Top:* sleep hypnogram highlighting the sleep states considered for analyses (NREM 2-3; grey rectangle). *Bottom:* multi-taper spectrogram (scalp EEG). (**b**) Example of ripple detection (accepted events: green; rejected events: red). *Top*: Unfiltered iEEG signal. *Middle:* Band-pass filtered signal (80-120 Hz). *Bottom:* Peak detection of ripple candidates (colored traces) on the normalized signal envelope. (**c**) Representative events. Inclusion criteria differentiate SW-R (left column) from rejected high-frequency bursts (middle column) and epileptic activity (right column). (**d**) Single-subject MTL SW-R. *Left:* Average waveform (mean ± SEM). *Middle:* Power spectrum relative to the ripple trough (± .1 s; black trace) highlighting a distinct peak (arrow) above the 1/f component (colored trace). *Right:* Time-frequency spectrogram highlights the transient, frequency-specific power increase in the ripple band. (**e**) Single-subject PFC SW-R. Same conventions as in panel **d**. (**f**) Group-level (*N* = 14) electrode coverage in MNI space of intracranial electrodes in the MTL (*N* = 82; red) and PFC (*N* = 339; blue). (**g**) Group-level MTL SW-R (*N* = 55,840). *Left:* Average ripple waveform (red) and superimposed sharp wave (<2 Hz; black). Insets highlight zoomed SW-R (*Left*, ± .1s) and nesting within the sharp-wave trough (R*ight*). *Top and Bottom Right:* Spectral SW-R representations; same conventions as in panel **d**. (**h**) Group-level PFC SW-R (*N* = 259,688). Same conventions as in panel **g**. (**i**) Ripple characteristics and their topographical depiction for MTL and PFC channels. From *Left* to *Right:* distributions of ripple frequency, amplitude, duration, and density (probability density functions; box plots represent median, 1^st^/3^rd^ quartiles and extreme values).

### Uniform morphology of MTL and PFC ripples

Having established the presence of SW-R in both the human MTL and PFC, we next directly compared their characteristics. MTL and PFC SW-R were indistinguishable in terms of peak frequency (*t*13 = −1.68, *p* = .1169, *d* = .45; **Fig. 1i** and ***SI Appendix*, Table S3**) and amplitude (*t* 13 = −1.69, *p* = .1153, *d* = .45), whereas PFC ripples exhibited a moderately longer duration (*t* 13 = −3.44, *p* = .0044, *d* = .92) and a slightly higher density (*t*13 = −3.07, *p* = .0089, *d* = .80) as compared to MTL ripples. Moreover, morphological similarity was evaluated by correlating the SW-R between a representative seed channel per region (i.e., channel with the strongest correlation to the 1^st^ principal component of the channel-averaged, ripple-locked data; ripple trough ± .05 s; *Methods*) with all other electrodes, either *within-* or *between-regions*. This analysis revealed that ripple morphology was highly consistent *within every region* (MTL: *r*^2^ = .51 ± .03; PFC: *r*^2^ = .42 ± .02; mean ± SEM), but also *between regions* (MTL: *r*^2^ = .44 ± .03; PFC: *r*^2^ = .46 ± .05) as no significant differences for *within-* vs. *between*-region comparisons were observed (MTL: *t*13 = 1.90, *p* = .0800, *d* = .51; PFC: *t*13 = .97, *p* = .3499, *d* = .26). Collectively, these findings demonstrate that SW-R reflect an inherent feature of human sleep and constitute a characteristic electrophysiological phenomenon in both the human MTL as well as the PFC.

### Cortical states shape sharp-wave ripple expression and timing

Previous work in rodents indicated that cortical up- and down-down states (as indexed by SO phase) modulate the expression of hippocampal SW-R (Mölle et al., 2006). Hence, we subsequently determined how SO shape the expression and precise temporal relationship between MTL and PFC SW-R (**Fig. 2a**). SW-R expression was highly heterogenous over the course of a SO in both the MTL and PFC (**Fig. 2b**), with significantly increased ripple rate during the up-state (MTL: *p*_uncorrected_ ≤ .0489, from −1.79 to −1.14 s; PFC: *p* = .0035, *d* = 1.23; from −1.99 to −1.14 s; cluster test), and suppression during the down-state (SO trough; MTL: *p* = .0006, *d* = 1.09, from .03 to 1.14 s; PFC: *p* ≤ .0001, *d* = 3.84, from −.53 to .33 s). Next, we determined whether MTL ripples preceded PFC ripples over the course of a SO. Although SW-R peaks relative to the SO trough were on average earlier for the MTL (−1.77 ± .15 s, mean ± SEM) as compared to PFC (−1.48 ± .12 s), no significant difference was observed (**Fig. 2c**; *t*_13_ = −1.44, *p* = .1745, *d* = .38), which directly results from the overall heterogenous SW-R timing in both regions (cf. **Fig. 2a**).

**Fig. 2.**
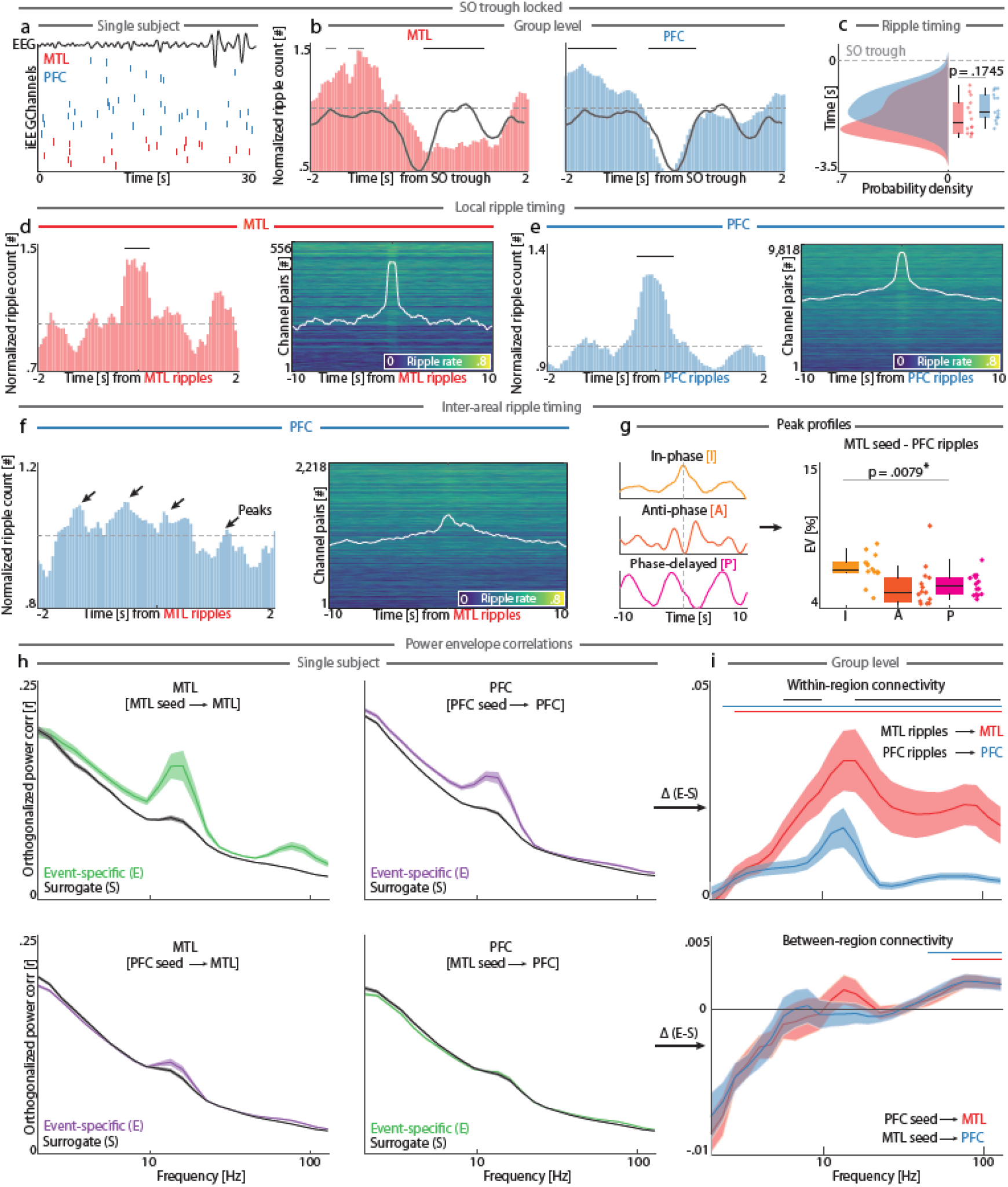
Temporal coordination of SW-R. (**a**) Single-subject SW-R raster (MTL: red; PFC: blue) relative to SO activity (scalp EEG, black). Note the grouping of MTL ripples. (**b**) *Left:* Histogram of MTL ripples relative to the SO trough highlights a non-uniform distribution across the SO cycle. *Right:* Histogram of PFC ripples relative to the SO trough. (**c**) Ripple timing relative to the SO down-state. Box plots represent median, 1^st^/3^rd^ quartiles and extreme values. Dashed horizontal line represents SO trough. (**d**) Within-region timing of MTL SW-R (*Methods). Left:* Normalized histogram of average SW-R rate across channels relative to SW-R on a representative MTL seed. *Right:* Stacked averages for all channel pairs. (**e**) Within-region timing of PFC SW-R. Same conventions as panels in **d**. (**f**) Inter-areal timing of PFC SW-R relative to MTL SW-R. Same convention as panels **d** and **e**. Note the multiple distinct peaks of the distribution (left, black arrows). (**g**) *Between-region* ripple peak profiles. *Left:* The most prevalent SW-R co-occurrence peak profiles. *Middle:* Distribution of peak profiles, with the *in-phase* profile being more prevalent as compared to the *anti-phase* or *phase-delayed* profile. (**h**) Single-subject orthogonalized power correlations (mean ± SEM) illustrate *within-region (Top*) as well as *inter-areal (Bottom*) event-specific connectivity upon SW-R (colored traces) relative to surrogate distributions (black traces). (**i**) Group-level differences between event-locked and surrogate distributions. *Top*: Increased *within-region* connectivity upon SW-R across several canonical frequency-bands. Colored lines indicate statistical significance (red/blue relative to 0, black condition difference). *Bottom: Inter-areal* connectivity relative to MTL (blue) and PFC (red) seeds. Same conventions as in panel **h**.

In order to quantify SW-R expression *within* a given region, we next determined their co-occurrence (*Methods*). We observed that ripples preferentially occurred simultaneously in multiple channels in the MTL (**Fig. 2d**; *p* = .0080, *d* = .82, from −.28 to .23 s; cluster test) and PFC (**Fig. 2e;** *p* = .0003, *d* = 1.24, from −.38 to .23 s). This set of findings demonstrates that *within-region* SW-R are temporally coordinated. Subsequently, we also assessed how SW-R were coordinated *between-regions*. For MTL-PFC SW-R coordination, we observed a more heterogeneous, albeit temporally-structured, profile as illustrated by multiple distinct peaks across time (**Fig. 2f**). To further describe this heterogeneous coordination profile, we computed principal component analysis (PCA) and identified three cardinal profiles from its first 10 principal components (**Fig. 2g**; in-phase, *anti-phase* and *phase-delayed*; each phase profile quantified by percent explained variance; %EV) that capture the relative timing of coordinated SW-R expression between the MTL and PFC. We observed a main effect of factor *profile* (*F*_2,39_ = 5.49, *p* = .0079, *η^2^* = .18; RM-ANOVA), with post-hoc tests revealing that the *in-phase* profile was more prevalent (7.03 ± .29%, mean ± SEM) as compared to *anti-phase* (5.68 ± .42%, *t*_13_ = 4.08, *p* = .0013, *d* = 1.09) and the *phase-delayed* profiles (5.48 ± .22%, *t*_13_ = 3.61, *p* = .0032, *d* = .96) separately. Notably, the *in-phase* profile was less prevalent than the combination of the other profiles (*t*_13_ = −9.70, *p* < .0001, *d* = 2.59). In sum, these observations establish that only a fraction of all SW-R occur simultaneously in the MTL and PFC, and hence raise the question how SW-R coordination between regions may facilitate the hippocampal-neocortical dialogue.

### SW-R modulate functional connectivity in the MTL-PFC network

To further quantify if MTL-PFC coordinated SW-R contribute to between-region information transfer, we next assessed if SW-R are functionally coupled. Employing functional connectivity analyses that rigorously account for field-spread in the cortical tissue (Hipp et al., 2012), we determined both *within-* and *between-region* coupling. Already on the single subject level (**Fig. 2h**), we observed ripple-mediated, band-limited functional connectivity that included the spindle-(12-16 Hz) and ripple-bands (80-120 Hz). For group-level analyses, we first accounted for a possible contribution of oscillatory power to the connectivity estimates by obtaining a power-matched surrogate distribution (*Methods*), which we subsequently subtracted from the observed estimates (**Fig. 2i**). This approach effectively controls for increased power during SW-R events.

We observed increased functional connectivity across multiple canonical frequency bands with distinct peaks in the spindle- and ripple-bands, both *within* the MTL (**Fig. 2i**; *p* < .0001, *d* = 1.32; cluster test) and *within* the PFC (*p* < .0001, *d* = 1.95; cluster test); mirroring the SW-R co-occurrence (cf. **Fig. 2d and e**). Together, these observations highlight the fact that cardinal sleep oscillations, such as SO (cf. **Fig. 2b**) and spindles (peaks in **Fig. 2i**; MTL: 16 Hz; PFC: 13.5 Hz) shape SW-R expression. Intriguingly, *within-region* connectivity was stronger in the MTL compared to the PFC (1^st^ cluster: 16-128 Hz, *p* = .0028 *d* = .89; 2^nd^ cluster: 5.66 to 9.51 Hz; *p* = .045, *d* = .83; cluster tests).

In contrast to *within-region* functional connectivity, MTL-PFC connectivity during the SW-R was largely confined to the ripple-band (**Fig. 2i**; MTL seed: *p* = .0058, *d* = .92, from 45.25-128 Hz; PFC seed: *p* < .0001, *d* = 1.17, from 64-128 Hz; cluster test). We did not observe significant differences between MTL- and PFC-seeded connectivity estimates, albeit a trend in the spindle frequency range (all *p_uncorrected_* ≥ .0805), indicating that PFC-ripple-mediated connectivity is accompanied by synchronized spindle activity, in line with previous findings (Helfrich et al., 2019; Staresina et al., 2015). In sum, these observations demonstrate coordinated SW-R activity between the MTL and PFC, possibly to temporally structure population activity in support of inter-areal information transfer.

### MTL and PFC SW-R differentially shape population activity

Having established that SW-R activity coordinates ripple expression in both regions as well as their functional interactions, these observations raise the question how SW-R exert their influence on *local* and *distant* neuronal populations in support of memory formation. A testable hypothesis arising from our observations is that SW-R should trigger state transitions in both the PFC and MTL to optimize inter-areal information reactivation and transfer. Here, we estimated high-frequency activity (120-200 Hz; HFA) as a surrogate marker of multi-unit population activity (Gallego-Carracedo et al., 2022; Leszczynski et al., 2020; Ray & Maunsell, 2011; Rich & Wallis, 2017). Note that we excluded the human ripple-band (80-120 Hz) from our HFA definition to avoid confounded estimates (***SI Appendix,*** **Fig. S1**).

Since our aforementioned *within-region* co-occurrence (**Fig. 2d and e**) and connectivity analyses (cf. **Fig. 2i**) demonstrated that MTL SW-R strongly synchronize MTL activity, we first tested whether MTL SW-R also trigger a stereotypical, low-dimensional population response. Therefore, we estimated the dimensionality of the population response from the eigenspectra as defined by PCA. Dimensionality can be inferred from the shape of the eigenspectrum (Gallego et al., 2017), where low-dimensional representations (i.e., the first few dimensions explain most variance) exhibit a steep decay function, while high-dimensional representations are characterized by a shallow decay function (**Fig. 3a** and *Methods;* note regional differences in number of electrodes was stratified through permutation). As predicted, population responses were lower-dimensional in the MTL compared to the PFC upon *local* SW-R occurrence (**Fig. 3b**; *t*_11_ = −3.56, *p* = .0045, *d* = 1.03; note that two participants with few MTL electrodes (*N* < 3) were excluded to reliably estimate population eigenspectra). Critically, the same pattern was observed for SW-R that emerged in the other region; i.e., we assessed the MTL population activity upon *distant* PFC ripples and vice versa (**Fig. 3b**; *t*_11_ = −4.23, *p* = .0014, *d* = 1.22). These results support the idea that the overall SW-R-mediated MTL population response is highly stereotypical and therefore low-dimensional, whereas the PFC response exhibits variable and rich spatiotemporal patterns.

**Fig. 3.**
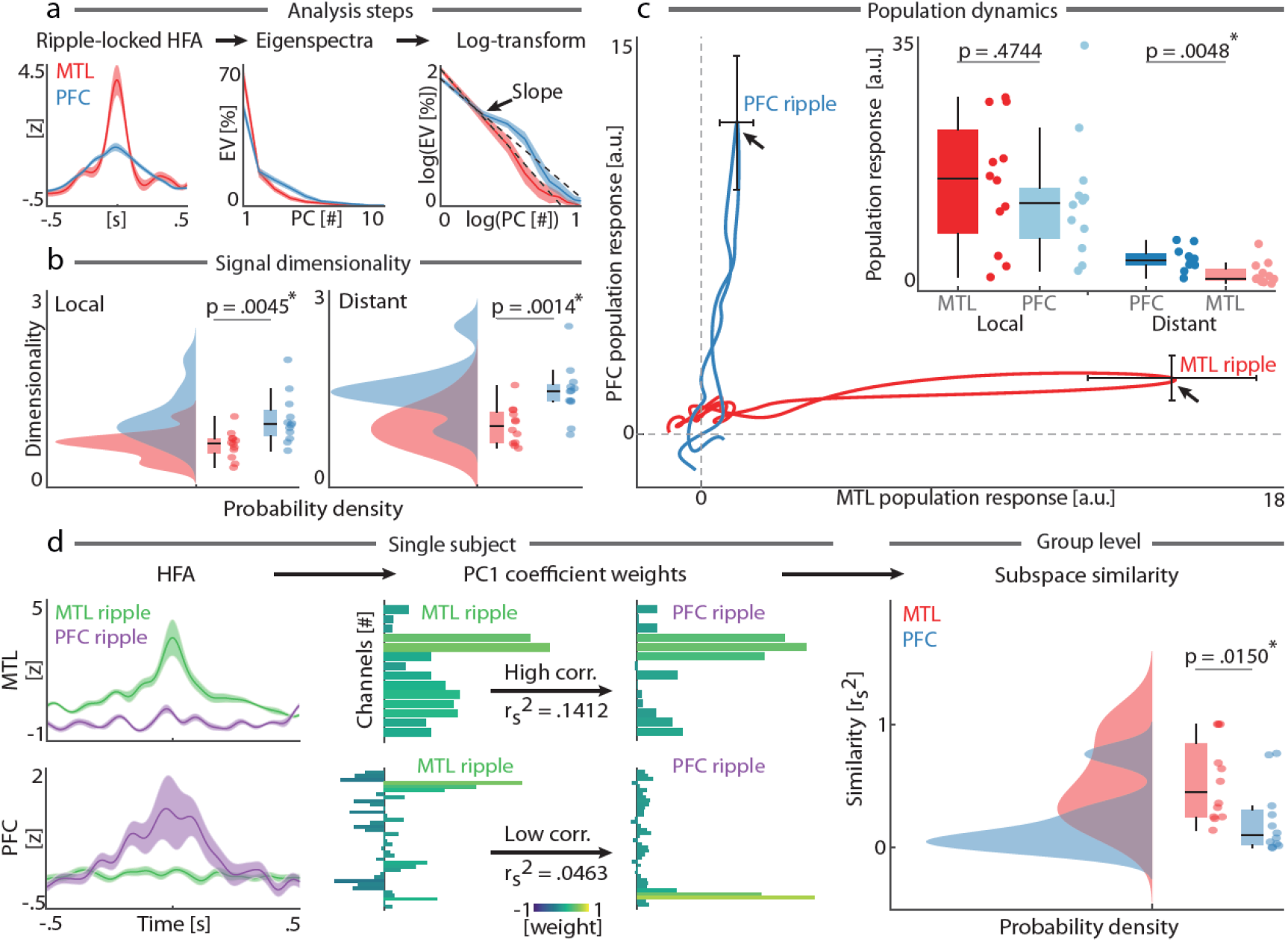
SW-R mediate MTL and PFC population activity. (**a**) Quantification of dimensionality upon SW-R. *Left:* Mean HFA time-locked to local ripples. *Middle:* Population eigenspectra upon SW-R (linear axis). *Right:* The decay of the eigenspectrum in log-log-spacing (black dotted line) indexes dimensionality. A steeper decay reflects lower dimensionality. (**b**) Group-level dimensionality of the population response upon SW-R is higher for the PFC than the MTL (relative to both *local* and *distant* SW-R). Data are displayed as probability density functions. Box plots represent median, 1 ^st^/3^rd^ quartiles and extreme values. (**c**) Projection of ripple-locked population activity into a common 2D state space. Trajectories indicate population dynamics within the MTL-PFC network over time (± .5 s relative to ripple trough; black arrows; mean ± SEM). Inset shows that population responses did not differ between the MTL and PFC upon *local* SW-R, but showed a stronger population response of the PFC upon *distant* ripples; thus, indicating that MTL-PFC interactions are primarily driven by the MTL. (**d**) Subspace similarity. From *Left* to *Right*: ripple-locked HFA dominant population responses, channel weights for the 1^st^ principal component (note the high similarity for the MTL as compared to the PFC), and group-level correlations per region. Subspace similarity between *local* and *distant* ripples was higher for the MTL as compared to the PFC.

We subsequently visualized and quantified the population trajectories upon SW-R by projecting the dominant ripple-locked population response (first principal component of ripple-locked HFA per region) into 2D state-space (**Fig. 3c**; *Methods*). This analysis revealed that SW-R-triggered population trajectories mainly transverse along the main population axis, i.e. MTL SW-R exert their main effect on the MTL population, whereas PFC SW-R mainly drive the PFC response. The influence of SW-R on the *local population* did not differ between the MTL and PFC (**Fig. 3c**, *Inset Left; t11* = .741, *p* = .4744, *d* = .21). In addition to driving the local population activity, SW-R also modulate the *distant* population activity. Here, we observed a clear dissociation, with MTL SW-R exhibiting a significantly stronger influence on PFC population activity than vice versa (**Fig. 3c**, *Inset Right*; *t*_11_ = 3.522, *p* = .0048, *d* = 1.02). In sum, these results demonstrate that MTL SW-R exert a strong and directed influence on the *temporal dynamics* of both *local* MTL as well as *distant* PFC population activity.

Having characterized the population trajectories over time, we next determined how SW-R modulate the network configuration, i.e. the *spatial dynamics*. Therefore, we first extracted the channel weight contributions to the dominant population response. We then quantified the subspace similarity of every region, depending on whether the SW-R emerged *locally* or in the *distant* region (**Fig. 3d**; correlation of subspace weights). In both regions, we identified a subspace that was preferentially modulated by *local* as well as *inter-areal* ripples (as indicated by a subspace similarity > 0; MTL: *t*_11_ = 5.71, *p* = .0001, *d* = 1.65; PFC: *t*_11_ = 2.73, *p* = .0197, *d* = .79). We observed a higher subspace similarity for the MTL (*r*^2^ = .54 ± .09, mean ± SEM) as compared to the PFC (**Fig. 3d**, *Right*; *t*_11_ = 2.88, *p* = .0150, *d* = .83, *r*^2^ = .22 ± .08), suggesting that the MTL response is highly stereotypical, irrespective of whether the SW-R emerged in the MTL or PFC. In contrast, PFC activity was differentially modulated by MTL and PFC SW-R.

In sum, we provide converging evidence across multiple uni- and multivariate analyses that MTL population dynamics are inherently low-dimensional, whereas activity in the PFC is high-dimensional. This functional dissociation might provide the scaffold for directed information transfer from the MTL to PFC, where a synchronized MTL activity provides the necessary means to imprint new information onto neocortical circuits.

### SW-R prioritize directed information transfer from the MTL to PFC

Finally, we directly probed the inter-areal information transfer between MTL and PFC upon SW-R. In light of the systems consolidation theory and following our findings, we hypothesized that information transfer should selectively increase upon an MTL SW-R from the MTL to PFC. To test this idea, we obtained time-resolved mutual information (MI; **Fig. 4a**) between both network nodes. Undirectional MI showed increased shared information between MTL and PFC upon MTL ripples (**Fig. 4b**; *p* = .0264, *d* = .88, from 1.55 to 2 s; cluster test). In contrast, PFC ripples decreased shared information between the MTL and PFC (**Fig. 4b**; *p* = .0074, *d* = 1.77, from 0 to .7 s; cluster test). Subsequently, we resolved the direction of the information flow (*Methods*) and observed that information flows almost exclusively from the MTL to PFC (**Fig. 4c**), but not from the PFC to MTL (**Fig. 4d**). Specifically, MI upon MTL SW-R increased shared MTL-to-PFC information (**Fig. 4c**; *p* = .0120, *d* = .93, from 1.65 to 2 s; cluster test). Critically, PFC SW-R reduced directional information flow from the MTL to the PFC (**Fig. 4c**; *p* = .0092, *d* = 1.49, from 0 to .70 s). In contrast, MTL SW-R decreased shared PFC-to-MTL information (**Fig. 4d**; *p* = .0138, *d* = 1.04, from .40 to .75 s), whereas no effect was observed upon PFC SW-R (**Fig. 4d**; all *p* ≥ .1942).

**Fig. 4.**
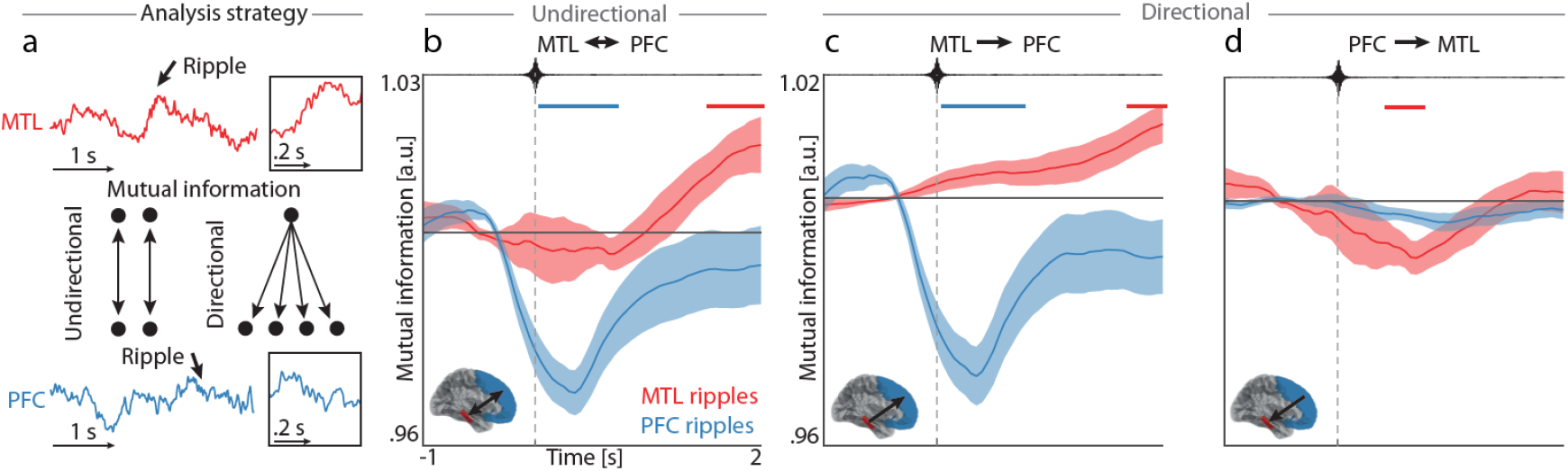
SW-R orchestrate directed MTL-PFC information transfer. (**a**) Analysis strategy to assess undirectional and directional mutual information (MI) upon SW-R. Box insets highlight ripple oscillations (± .1 s). (**b**) Inter-areal undirectional mutual information upon ripples (mean ± SEM). MTL SW-R increased mutual information, whereas PFC ripples impede communication. (**c**) Directionality analyses show information transfer from the MTL to PFC upon MTL SW-R, whereas input from the MTL is reduced upon PFC ripples. (**d**) Directional analyses from the PFC to the MTL using the same conventions as **c**. Note no increase in directional MI is observed in the MTL following either MTL or PFC SW-R.

Taken together, these results provide robust evidence that the SW-R is a key substrate for directed information transfer from the MTL to PFC, providing either the necessary means to transfer information (upon the MTL SW-R) or block information (upon PFC SW-R); thus, preventing interference of new input with the processing of recently received information.

### Functional dissociation between MTL and PFC ripples is species independent

Previously, SW-R were thought to reflect a MTL-specific electrophysiological signature, but have recently also been observed in neocortical areas. The majority of studies on SW-R focused on the rodent hippocampus, highlighting the presence of SW-R in the range from 110-200 Hz. In humans, putative homologues of the rodent SW-R are typically slower (80-120 Hz). To validate our approach and present findings in humans, we subsequently replicated all key univariate analyses utilizing intracranial recordings from male Long Evans rats (*N* = 4; statistics are reported on the pseudo-population level). Surface EEG and intracranial local-field potentials (LFP) were recorded from the MTL and PFC using two platinum electrodes **(Fig. 5a;** Oyanedel et al., 2020; see ***SI Appendix*** for details regarding surgery, recordings, histology, and sleep stage determination). Ripples were detected using the aforementioned algorithm, accounting for a wide frequency range (60-240 Hz; **Fig. 5b**). Inclusion criteria regarding frequency, amplitude, duration, and sharp transients were kept constant between human and rodent data.

**Fig. 5.**
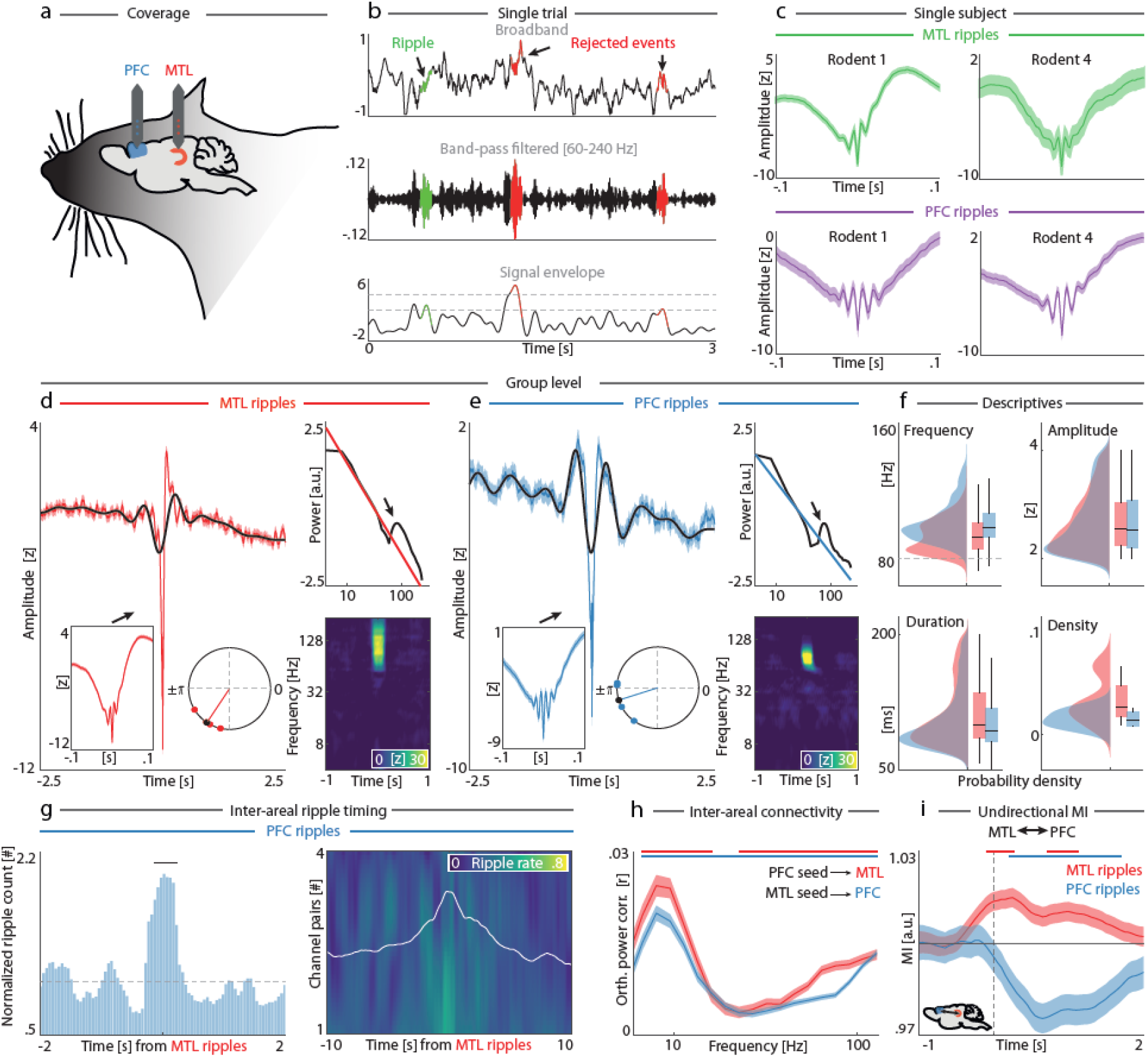
Replication of SW-R detection and key analyses in rodents. (**a**) Schematic of recording location in the rat, utilizing two platinum electrodes located in the medial prefrontal cortex (PFC) and dorsal hippocampus (MTL). (**b**) SW-R detection on rodent data utilized the same algorithm and inclusion criteria as previously applied in humans (60 to 240 Hz). (**c**) Single-subject MTL (*Top*) and PFC (*Bottom*) SW-R. (**d**) Group-level (*N*=4) MTL SW-R (*N* = 4,844). *Left:* Average ripple waveform (red) and superimposed sharp wave (black). Insets highlight zoomed SW-R (*Left*, ± .1 s) and nesting within the sharp-wave trough (*Right). Right:* Spectral SW-R representations. (**e**) Group-level PFC SW-R (*N* = 2,359). Same conventions as in panel **d**. (**f**) Ripple characteristics of all detected events. Horizontal dashed line indicates the 90 Hz ripple peak observed in humans (cf. **Fig. 1*I*; *SI Appendix*, Fig. S2c**). Data are displayed as probability density functions. Box plots represent median, 1^st^/3^rd^ quartiles, and extreme values. Group-level details on ripple morphology are provided in the Supplementary Materials (***SI Appendix,*** **Table S4**). (**g**) *Inter-areal* SW-R coordination. PFC ripples co-occurred in close temporal proximity relative to MTL ripples. (**h**) Connectivity between the MTL and PFC upon SW-R. (**i**) Mutual information (MI) between the MTL and PFC increased upon MTL ripples (red), whereas PFC ripples inhibited inter-areal communication (blue).

Rodent SW-R showed highly comparable morphological characteristics as observed in humans (single-subject examples shown in **Fig. 5c**). First, SW-R frequencies were faster than in humans, both in the MTL (99.96 ± .18 Hz, mean ± SEM) and PFC (105.62 ± .35 Hz). Second, ripple nesting within the sharp-wave trough was present in both the MTL (**Fig. 5d;** *p* < .0001, v = 445.4, V-test against π; coupling direction 231.4° ± 1.1°, mean ± SEM) and PFC (**Fig. 5e;** *p* < .0001, v = 335.6, V-test against π; coupling direction 197.3° ± 1.1°). Third, ripples were oscillatory in nature, resulting in a frequency-specific, transient power increase relative to the 1/f× background activity (**Fig. 5d and e**, *Right*). Fourth, SW-R morphology (**Fig. 5f**) between the MTL and PFC was indistinguishable in terms of amplitude (*t*_7201_ = .06, *p* = .9498, *d* = .00; across events) and density (*t*6 = 1.29, *p* = .2437, *d* = .91; across channels), albeit with a lower frequency for the MTL as compared to the PFC (*t*_7201_ = −15.52, *p* < .0001, *d* = .39) and slightly longer duration for the MTL (.112 ± .001 s, mean ± SEM) as compared to the PFC (*t*_7201_ = 8.89, *p* < .0001, *d* = .22, .105 ± .001 s). These results confirm that MTL and PFC SW-R exhibit a near-uniform morphology that reflects an inherent signature in both species.

Next, we also assessed the inter-areal temporal coordination and functional connectivity relative to SW-R. Note that comparisons in rodents were confined to uni- or bivariate comparisons, given that only one recording electrode was available per region. Confirming previous results in rodents (Khodagholy et al., 2017), but in contrast to our results in humans, we observed that PFC ripples occurred in close temporal proximity with MTL ripples (**Fig. 5g**; *p* < .0001, *d* = 2.52, from − .23 to .23 s, cluster test). Yet, mimicking the human results (cf. **Fig. 2i**), inter-areal connectivity was modulated by SW-R from the MTL (**Fig. 5h**; 1^st^ cluster: 23-128 Hz, *p* = .0001 *d* = .45; 2^nd^ cluster: 4 to 16 Hz; *p* = .0001, *d* = .45; cluster tests) and PFC (*p* = .0001 *d* = .57, from 5.7 to 128 Hz). Lastly, ripple-locked communication between the MTL and PFC was assessed through their change in mutual information. Again, we observed that MI increased between the MTL and PFC upon MTL SW-R (**Fig. 5i**; 1^st^ cluster: from −.1 to .25 s, *p* = .0012, *d* = .13; 2^nd^ cluster: from .7 to 1.1 s; *p* = .0096, *d* = .07; cluster tests), whereas a significant drop in shared information was observed upon PFC ripples (*p* = .0004, *d* = .15, from .2 to 1.7 s). Replicating our findings in humans, inter-areal communication between MTL and PFC in rodents was facilitated by MTL ripples, whereas communication was impeded by PFC ripples.

## Discussion

Our results reveal a striking dissociation between MTL and PFC SW-R in support of hippocampal-neocortical information processing in both human and rodent cortex. SW-R exhibit a near uniform morphology in both regions, but fulfil distinct functional roles that jointly support a timed information hand-off from the hippocampus to the neocortex. While MTL SW-R synchronize local and global networks to trigger directional hippocampal-to-neocortical information flow, cortical SW-R diversify the local population response and block subsequent input from the MTL. These results describe a division-of-labor between MTL (sender) and PFC (receiver), with SW-R constituting a key substrate that structures hippocampal-neocortical dynamics in space and time. In sum, converging evidence across species and brain regions demonstrates that coordinated SW-R activity provides temporal reference frames that segregate hippocampal information reactivation and subsequent neocortical processing.

### Uniform SW-R morphology across regions and species

SW-R hallmark hippocampal dynamics during sleep. The majority of studies focused on the rodent hippocampus during sleep, but SW-R have also been described in hippocampal and adjacent MTL structures in humans using intracranial recordings (Axmacher et al., 2008; Bragin et al., 1999; Chen et al., 2021; Helfrich et al., 2019; Norman et al., 2019; Staresina et al., 2015; Vaz et al., 2019). Recently, it became evident that SW-R are not exclusive to the MTL, but are also an inherent signature of neocortical circuits (Dickey, Verzhbinsky, Jiang, Rosen, Kajfez, Eskandar, et al., 2022; Khodagholy et al., 2017; Vaz et al., 2019). However, given that intracranial human recordings are typically obtained from epileptic patients, it has been debated whether the putative human homologues of SW-R reflect true ripples or whether they constitute artifacts (Le Van Quyen et al., 2010). Moreover, SW-R are typically viewed as a unitary phenomenon (Buzsaki, 2015), but emerging evidence suggests that distinct SW-R features, such as duration (Fernandez-Ruiz et al., 2019) or density (Norman et al., 2019) index functional specialization. If similar considerations apply to neocortical SW-R in the human brain (Ngo et al., 2020) is currently not known. The paucity of knowledge is a direct consequence of the lack of comparative evidence linking rodent and human SW-R.

Here, we confirm that SW-R are an inherent electrophysiological phenomenon of the sleeping brain. Critically, we observe highly comparable SW-R in the MTL and PFC for humans and rodents. Validating our detection approach in rodents that did not suffer from epilepsy indicates that true SW-R were successfully disentangled from pathologic activity in human intracranial recordings. In line with previous reports (Bragin et al., 1999; Buzsaki, 2015; Jiang et al., 2020; Oyanedel et al., 2020), we observed a higher frequency for ripples in rodents (~100 Hz) than in humans (~ 90 Hz), while the majority of morphological features, including coupling to the sharp-wave trough, remained highly consistent. In sum, our findings establish that SW-R are in fact a widespread cortical phenomenon and demonstrate the feasibility of between-region and -species comparisons.

### SW-R structure information reactivation, transfer and processing in space and time

The demonstration that SW-R are ubiquitous raises the question how they exert their influence on large-scale cortical networks to support memory reactivation, transfer, and consolidation. Hippocampal SW-R have long been recognized as the most synchronous population pattern of the mammalian brain that modulates both local as well as network activity (Buzsaki, 2015). SW-R also structure replay (Olafsdottir et al., 2018), i.e. the reactivation and recapitulation of firing patterns that were first present during encoding, which subsequently become strengthened: thereby, constituting a key signature of the consolidation process. At the network level, SW-R have been shown to modulate activity at distant cortical sites (Logothetis et al., 2012; Skelin et al., 2021) and are embedded in a precisely-timed oscillatory hierarchy of cardinal sleep oscillations, including neocortical SO and thalamocortical spindles (Born & Wilhelm, 2012; Clemens et al., 2007; Staresina et al., 2015). While MTL and neocortical SW-R might be coupled (Dickey, Verzhbinsky, Jiang, Rosen, Kajfez, Stedelin, et al., 2022; Khodagholy et al., 2017; Vaz et al., 2019), little is known about their specific function.

Our results now demonstrate that the human MTL SW-R strongly synchronize population activity, both locally in the MTL as well as in the PFC. Regarding the *within-MTL* dynamics, we demonstrate that MTL SW-R (i) are temporally coherent (**Fig. 2d**); (ii) synchronize population activity in multiple canonical frequency bands, most pronounced in the spindle- and ripple-bands (**Fig. 2i**); thereby, (iii) promoting a stereotypical, low-dimensional population response (**Fig. 3b**) that (iv) establishes MTL-PFC functional connectivity in the ripple-band (**Fig. 2i**) to (v) selectively channel MTL-dependent information to the PFC (**Fig. 4b and c**).

In stark contrast to its MTL counterpart, our analyses reveal distinct functional properties of the PFC SW-R, which are coherent within the PFC (**Fig. 2e**), but exhibit a more complex temporal relationship to the MTL SW-R as previously assumed (**Fig. 2f**). (i) Prefrontal SW-R only weakly synchronize local and inter-areal networks (**Fig. 2i** and **3c**); thus, (ii) diversifying the population response, which, as a direct consequence (iii) is high-dimensional and not confined to a low-dimensional subspace (**Fig. 3b-d**). The increased dimensionality of neocortical circuits might reflect a substrate that allows the imprinting of novel mnemonic patterns onto existing circuits in support of memory consolidation. In line with this consideration, we also observed that (iv) PFC SW-R decrease the directional information flow from the MTL to PFC (**Fig. 4b** and **c**), possibly reducing interference to enable sequential processing of information as provided upon an MTL ripple. Collectively, this set of findings demonstrates that the interplay of MTL and PFC ripples structures inter-areal information transfer in support of memory consolidation in space and time. These findings further suggest that SW-R are irrevocably linked to MTL-PFC directed network interactions, even if the respective SW-R occur outside the MTL, since virtually all information transfer occurs from the MTL to PFC and not vice versa (**Fig. 4d**). Lastly, our observations in rodents replicate and extend these findings in humans and further confirm that SW-R serve as endogenous timing mechanisms that (i) establish temporally- and frequency-specific communication channels (**Fig. 5g** and ***H***) to (ii) mediate MTL-to-PFC information transfer (**Fig. 5i**). Collectively, this set of findings provides converging evidence that sleep oscillations are evolutionary conserved, functional substrates that organize the transfer and processing of information to enable long-term memory consolidation.

## Materials and Methods

### Subjects

Patients diagnosed with pharmacoresistant epilepsy who underwent invasive monitoring with implanted electrodes in the MTL and PFC were recruited from the University of California Irvine Medical Center, USA (*N* = 14; *M_age_* = 36.79 ± 13.28 years, range 19-58 years; 9 female). Intracranial recordings were conducted as part of the pre-surgical invasive monitoring to identify the seizure onset zone. The intracranial depth electrodes (Ad-Tech) were implanted unilaterally (*n* = 1), bilaterally (*n* = 12), or unilaterally combined with grid electrodes (*n* = 1). Electrode localization was determined in native space by two independent neurologists, whereas group-level visualization of electrode location was done in normalized MNI space (**Fig. 1f**). Details concerning patient diagnosis, electrode number, lateralization, and localization are provided in the Supplementary Materials (***SI Appendix,*** **Table S1**). All subjects provided written informed consent prior to study participation. All protocols concerning data acquisition and analyses were approved by the Institutional Review Board at the University of California, Irvine (protocol number: 2014–1522) and the Committee for Protection of Human Subjects at the University of California, Berkeley (Protocol number: 2010-02-783). The study was conducted in accordance with the 6^th^ Declaration of Helsinki.

### Procedure

Data were simultaneously recorded from electrodes located in the MTL and PFC, as well as from additional scalp EEG electrodes (Fz, Cz, C3, C4, and Oz placed according to the international 10-20 system), together with electrooculogram (EOG; four electrodes placed around the left and right outer canthi) and electromyogram (EMG). All electrophysiological data were acquired using a Nihon Kohden recording system (model JE120A; 256-channels), were digitally sampled at 5000 Hz, and analog filtered at .01 Hz.

Full-night recordings started between 19:30 – 22:00 and lasted 10-12h (***SI Appendix*, Table S1**). One seizure-free night of sleep was analyzed per patient. Sleep scoring was conducted by an expert rater in accordance with established criteria (Iber et al., 2007); **Fig. 1a, *Top***) using scalp EEG as well as EOG and EMG. Average sleep duration was 447 ± 123 min (mean ± SD; range 181 to 649 min), of which 74% ± 8% was spent in sleep states NREM2-3 (range 64% to 86%). Single-subject details on sleep architecture are provided in the Supplementary Materials (***SI Appendix*, Table S2**).

### Data processing

Faulty electrodes, those associated with epileptiform activity, or those with a low ripple number (events < 100) were excluded from analyses. iEEG electrode locations were visually verified and cross-referenced with the AAL atlas (Tzourio-Mazoyer et al., 2002). For preprocessing, iEEG data were demeaned, detrended, rereferenced to a bipolar montage, and downsampled to 500 Hz. Similarly, EEG data were demeaned, detrended, rereferenced to a common scalp average, low-pass filtered at 50 Hz, and downsampled to 500 Hz. Analyses of scalp EEG focused on the singular electrode Fz (or channel Cz if Fz was unavailable). Data processing and analyses were carried out using Matlab R2021a (MathWorks Inc.). Preprocessing, segmentation, and time-frequency analyses utilized custom code in addition to functions from the FieldTrip toolbox (Oostenveld et al., 2011) and EEGLAB toolbox (Delorme & Makeig, 2004). Moreover, we used the FOOOF algorithm (“fitting oscillations & one-over f”; Donoghue et al., 2020) to assess the ripple-locked aperiodic components of the power spectra (**Fig. 1d, e,** and **h**).

### Event detection

#### Ripple detection

Ripples were detected following previously established criteria (Helfrich et al., 2019; Khodagholy et al., 2017; Vaz et al., 2019). Data were band-pass filtered between 80 and 120 Hz using an absolute Hilbert transform, smoothed using a 10 Hz low-pass filter, and z-normalized (**Fig. 1b**). Peaks in the high gamma signal that exceeded two standard deviations from the mean amplitude, had a duration between 25 and 200 ms, and a minimum peak distance of 500 ms were considered as candidate events. Events detected within the first and last minute of the recording were removed to avoid potential artifacts. All events were cross-referenced with sleep staging. Candidate events that met the following criteria were included in the analyses: (i) peak amplitude ≥ 2 and ≤ 4 (z-scored); (ii) frequency > 80 Hz; (iii) sharp transients (max peak amplitude differential) < 2 (z-scored); (iv) duration > median – SD; (v) occurrence during sleep states NREM 2-3. In case of overlap with other ripples ± 2.5 s relative to the ripple trough, we selected the ripple with the longest duration (Fernandez-Ruiz et al., 2019; Ngo et al., 2020). Finally, the raw iEEG signal was segmented in artifact-free epochs centered around the ripple troughs (± 2.5 s). These stringent selection criteria differentiated ripples from broadband bursts and pathologic activity (**Fig. 1c**; ***SI Appendix*, Fig. S1**). Mean ripple characteristics are reported per sleep state and region in the Supplementary Materials on group level (***SI Appendix*, Table S3**) as well as single-subject level (***SI Appendix*, Table S4**).

#### Slow wave detection

Slow waves were detected on scalp EEG following previously described criteria (Hahn et al., 2022; Hahn et al., 2020; Helfrich et al., 2019; Helfrich et al., 2018; Staresina et al., 2015). Data were high-pass filtered at .16 Hz and subsequently low-pass filtered at 1.25 Hz using a Butterworth IIR filter and filter orders of 2 and 6, respectively. Zero crossings were defined, after which only slow oscillations during NREM 3 with a duration between .8 and 2 s were considered for analyses.

### High-frequency activity

High-gamma activity (120 to 200 Hz; HFA) has been closely linked with multi-unit activity (Gallego-Carracedo et al., 2022; Leszczynski et al., 2020; Ray & Maunsell, 2011; Rich & Wallis, 2017) and was therefore used as a surrogate marker of population activity. Note that our frequency range for HFA is higher compared to previous studies (~80-150 Hz; c.f., Canolty et al., 2006; Ray & Maunsell, 2011; Rich & Wallis, 2017) to minimize a potential overlap with the ripple frequency band (80 to 120 Hz). HFA was determined using the following steps. First, the continuous, raw iEEG signal was bandstop filtered to account for the 60 Hz line noise and its harmonics (58 to 62 Hz; 118 to 122 Hz; & 178 to 182 Hz). Next, data were band-pass filtered per center frequency (120 to 200 Hz in 10 Hz steps; center frequency ± 5 Hz), followed by a Hilbert transform to determine the signal envelope, and z-normalization. Finally, we averaged over different sub-bands to obtain the mean HFA signal per electrode.

### Data analyses

#### Full-night spectrogram

A spectrogram of a full-night recording (**Fig. 1a, *Bottom***) was obtained following previously described analytical steps (Helfrich et al., 2019). Scalp EEG data were downsampled to 100 Hz, followed by z-normalization and segmentation into 30 s segments with a 95% overlap. Power spectra were computed per segment (frequency range .5 to 20 Hz in .5 Hz steps and ± .5 Hz frequency smoothing) using 29 Slepian tapers.

#### Ripple-locked event-related potential (ERP)

To illustrate ripple morphology, iEEG data were segmented per channel relative to their respective ripples per region (**Fig. 1d, e, g** and **h**; ripple trough ± 2.5 s).

#### Ripple-locked assessment of power spectrum densities and spectograms

For each electrode, we calculated the power spectrum density (ripple trough ± .1 s; Hanning taper with frequency range 4 to 120 Hz) and spectrogram (ripple trough ± 2.5 s; DPSS taper with frequency range 4 to 181 Hz in 45 logarithmical steps). The background components of the 1/f power spectra were estimated in log-log space from 4 to 120 Hz using the FOOOF algorithm (Donoghue et al., 2020). Region-specific spectra were determined by averaging over the associated electrodes. The aperiodic component was subsequently calculated on the averaged spectrum per region (**Fig. 1d, e, g,** and **h**).

#### Selection of seed electrode per region

To assess local and inter-areal interactions upon ripples, we identified a seed electrode whose ripples best represented the average ripple morphology per region per subject. First, iEEG data were segmented per channel relative to their individual ripples. The predominant ripple morphology was determined per region by conducting PCA on the channel-averaged ripple-locked data and extracting the time-series of the first principal component. The electrode with the highest correlation coefficient with the weights of the first principal component was identified as the representative seed electrode. Subsequent analyses were conducted time-locked to ripples per seed electrode.

#### Ripple waveform similarity

We assessed similarities in waveform shape by correlating the channel-averaged ripple time series (ripple trough ± .05 s) with the time series of the representative seed electrodes per region. Correlational values were subsequently squared and averaged per subject to yield one similarity score per subject. We subsequently contrasted similarity scores between *within-* and *between-region* ripples.

#### Ripple characteristics

We determined ripple characteristics (i.e., frequency, amplitude, and duration) per event and electrode (**Fig. 1 i**). Ripple density was determined per electrode by dividing the number of ripples by the time spent in the respective sleep states. Topographical representations of ripple descriptives show mean descriptive values per electrode.

#### Cluster-based permutation

We utilized Monte-Carlo cluster-based permutation tests (10,000 iterations) as implemented in Fieldtrip to correct for multiple comparisons. Clusters were determined in the time and frequency domains following two-tailed dependent t-tests and threshold at *p* < .05 (**Fig. 2b-e,** and **h**; **Fig. 4b-d, Fig. 5h** and **i**).

#### Cross-frequency coupling

Assessment of cross-frequency coupling between ripples and sharp waves (**Fig. 1g** and **h; Fig. 5d** and **e**) was conducted in accordance with the analysis steps of previous studies (Hahn et al., 2022; Hahn et al., 2020; Helfrich et al., 2019; Staresina et al., 2015). Sharp-wave phase angles of the Hilbert transform were determined upon ripple troughs following z-normalization and band-pass filtering (.3 to 2 Hz). Mean coupling direction and strength were calculated per channel per region using the Circular Statistics Toolbox (Berens, 2009). We subsequently determined mean circular direction and coupling strength per region per subject.

#### SO and ripple trough-locked peri-event histograms

Following SO detection on the EEG data, their co-occurrence with ripples was assessed for each iEEG electrode (SO trough ± 2.5 s; **Fig. 2b**). Peri-event histograms were subsequently smoothed and normalized to the grand mean per channel. Similarly, local and inter-areal ripple co-occurrence were assessed per region relative to ripples on representative seed electrodes (ripple trough ± 2.5 s; **Fig. 2d-f**). In addition, we assessed ripple co-occurrence for all channel pairs (ripple trough ± 10 s; **Fig. 2d-f**), sorted pairs according to ripple rate, and color-coded ripple rate over time.

#### Ripple peak profiles

The analysis of inter-areal ripple co-occurrence (**Fig. 2g**) showed higher variability in peak profiles as compared to local ripple co-occurrence (**Fig. 2d** and **e**). We therefore assessed the prominence of different peak profiles. Following the assessment of ripple co-occurrence for all channel pairs (**Fig. 2e,** *Right*), we identified the 10 most prominent peak profiles using PCA on group level. These profiles were subsequently categorized as *in-phase*, *anti-phase*, and *phase-delayed* (**Fig. 2g,** *Left*). This was repeated on single-subject level per region; now with the 10 most prominent peak profiles categorized based on their correlation with the group-level profiles. Peak profile prominence was defined as the average explained variance per category (**Fig. 2g,** *Right*).

#### Event-locked connectivity analyses

Connectivity between electrodes and regions was assessed relative to ripples occurring on representative seed electrodes per region of interest (ROI) per subject (**Fig. 2h**). Orthogonalized power correlations were calculated in accordance with previous studies (Helfrich et al., 2019; Hipp et al., 2012) for center frequencies from 2 to 128 Hz in 25 logarithmical steps. Continuous data were preprocessed by applying a band-pass filter per seed frequency (± ¼ center frequency) followed by segmentation relative to ripples on the seed electrode (ripple trough ± 2.5 s). We subsequently extracted the complex signal using the Hilbert transform. Next, we calculated the absolute orthogonalized power correlations per frequency pair, electrode, and subject between the two orthogonalized signals A and B (Eq. 1–2).

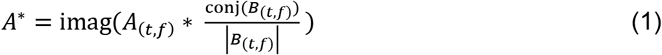

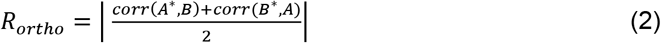

With A and B representing the phase series for the target and seed electrodes, respectively. Next, we compared the connectivity estimates calculated per ripple to a surrogate distribution. Here, we calculated orthogonalized power correlations between seed data for event N and target data for event N+1 (note that the last events were consequently discarded). The contrast between event-specific and surrogate distributions consequently shows the modulation of connectivity across frequencies upon ripples (**Fig. 2i; Fig. 5h**).

#### Signal dimensionality of the population response upon ripples

Dimensionality of the population response was assessed using the slope of the PCA eigenspectrum in log-log space. When the population response is highly stereotypical and of low dimensionality, the eigenspectrum shows a high offset (i.e., high explained variance of the first principal component) and a steep slope (i.e., the first few components predominantly explain the signal variance, whereas subsequent components make little contributions). Consequently, the slope of the eigenspectrum illustrates the dimensionality of the underlying signal.

We first segmented the HFA traces relative to ripples (*local* ripples - i.e., population response relative to ripples originating in the same region; *inter-areal* ripples - i.e., population response relative to ripples originating in the alternate region; **Fig. 3a,** *Left*). Next, the mean eigenspectra were calculated per region per subject through permutation; matching the number of electrodes between regions prior to conducting PCA over 1000 iterations (**Fig. 3a,** *Middle*). We subsequently determined the slopes of the PCA eigenspectra in log-log space (**Fig. 3a,** *Right*) by fitting a first-degree polynomial. Finally, dimensionality was expressed as the absolute of the slope.

#### Population dynamics upon ripples

We show population dynamics of MTL and PFC upon ripples originating in either region. For this step HFA data were utilized as a proxy of neuronal population firing (Ray & Maunsell, 2011; Rich & Wallis, 2017). First, representative seed electrodes were determined per ROI. HFA data were subsequently segmented relative to ripples on the seed electrode (**Fig. 3a,** *Left*). Second, we determined the predominant population response per region per subject by conducting PCA on the channel-averaged, ripple-locked HFA data and extracting the time series of the first principal component. Third, we averaged over subjects to determine the mean state-space population responses (ripple trough ± .5s; **Fig. 3c**). This state-space trajectory thus illustrates the population dynamics between MTL and PFC upon ripples over time. *Population response upon ripple troughs* was assessed by determining the Euclidean distance in high-dimensional space (maximum number of *N*_PCs_) for the MTL and PFC relative to surrogate distributions of zeros (**Fig. 3c,** *Inset*).

#### Subspace similarity

The similarity in population responses was assessed between *local* and *inter-areal* ripples. First, the HFA population response was determined relative to ripple (**Fig. 3d,** *Left*). Next, we conducted PCA on the ripple-locked HFA data and extracted the channel coefficients for PC1 (weights). These weights illustrate channel contributions to the dominant population response (**Fig. 3d,** *Middle*). We then correlated the channel weights upon local and inter-areal ripples per patient. High correlational values are indicative of high subspace similarity, meaning that the overall population response is similar between events. Correlational values were subsequently contrasted between regions (**Fig. 3d,** *Right*).

#### Mutual Information

We evaluated how ripples affect the information flow between the MTL and PFC by calculating event-locked directional and undirectional mutual information (MI). Analyses were conducted between a representative seed electrode per region and all electrodes of the target region (**Fig. 4a**). Target data were specified as all remaining available channels for both regions. The subsequent steps were done in accordance with previous studies (Helfrich et al., 2019; Quiroga & Panzeri, 2009). Seed and target data were segmented relative to ripples occurring on the seed electrode (ripple trough −1 to +2.25 s). The calculation of MI between seed & target data utilized a 400ms sliding window and 50ms step size. Data were binned per time window (*N*bins = 8; uniform bin count) and MI was calculated per time window, trial, channel pair, and subject (Eq. 3). MI traces were mean-normalized relative to baseline (−1 to 0 s relative to ripple trough). Undirectional MI was assessed between seed and target data with a uniform sliding window for both time series (**Fig. 4b, Fig. 5i**). In contrast, directional MI was assessed between a fixed window on the ripple (ripple trough ± .2 s) for a single time series relative to all time windows in the other time series (**Fig. 4c** and **d**). MI between time series X and Y was defined as:

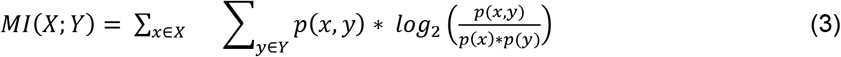

With *p*(*x,y*) being the joint probability function and *p*(*x*) and *p*(*y*) the class probabilities. MI traces were averaged over trials per channel and mean-normalized relative to the individual baseline (−1 to 0s). Topographical representations of MI (**Fig. 4c**) show the change in MI relative to the baseline per channel.

## Acknowledgments

This study was supported by the German Research Foundation, Emmy Noether program (DFG HE8329/2-1; RFH), Jung Foundation for Science and Research (RFH), Hertie Foundation (Network for Excellence in Clinical Neuroscience; RFH), and the Baden Wuerttemberg Foundation (Postdoc Fellowship; RFH).

## Author Contributions

F.J.V.S. and R.F.H. designed the study. J.D.L., J.J.L. and R.F.H. acquired and preprocessed the human data. M.A.H. and M. I. acquired and preprocessed the rodent data. F.J.V.S., M.A.H., J.D.L. and R.F.H. curated the data. F.J.V.S., J.W., and R.F.H. conducted statistical analyses. F.J.V.S. and R.F.H. drafted the manuscript. R.F.H. supervised the project. All authors commented on and edited the manuscript draft.

## Data and code availability

All data and custom code used for analyses are available from the corresponding author upon reasonable request.

## Additional information

**Correspondence** and requests for materials should be addressed to F.J.V.S or R.F.H.

