## Supplementary Materials for "An evolutionary conserved division-of-labor between hippocampal and neocortical sharp-wave ripples organizes information transfer during sleep"

**This file contains:**

Supplementary text

Supplementary figures S1-2

Supplementary tables S1-5

Supplementary Information references

**Human data**

**Co-occurrence between ripples and pathologic activity.** Electrodes that were associated with epileptiform activity (as determined by an epileptologist) were excluded from analyses. Yet, pathologic interictal epileptiform discharges (IEDs) could still sporadically occur on the remaining electrodes. We therefore evaluated the overlap between ripple events and IEDs relative to the ripple trough (± .05 s). Average co-occurrence per channel was low ($N_{channels}$ = 421; $N_{ripples}$ = 315,528; $N_{IEDs}$ = 480; Overlap = .15 ± .33% (mean ± SD); range = 0 to 3.04%). We consequently conclude that our detection and cleaning pipeline successfully distinguished between physiologic ripple and pathologic IED events.

**Rodent data**

**Animals.** The recordings were performed in four male Long Evans rats (Janvier, Le Genest-Saint-Isle, France, 280–340 g, 14–18 weeks old). Animals were kept on a 12 hr/12 hr light/dark cycle with lights off at 19:00 hr. Water and food were available ad libitum. All experimental procedures were approved by the University of Tübingen and the local institutions in charge of animal welfare (Regierungspräsidium Tübingen). A subset of the animals had been used in a previous study (Duran et al., 2018). Note that one rodent from the original dataset (Oyanedel et al., 2020) could not be analyzed given insufficient signal quality of the prefrontal LFP signal.

**Surgery.** Animals were anesthetized with an intraperitoneal injection of fentanyl (0.005 mg/kg of body weight), midazlolam (2.0 mg/kg), and medetomidin (0.15 mg/kg). They were placed into a stereotaxic frame and were supplemented with isoflurane (0.5%) if necessary. The scalp was exposed and five holes were drilled into the skull. Three EEG screw electrodes were implanted: one frontal electrode (AP: +2.6 mm, ML: −1.5 mm, with reference to Bregma), one parietal electrode (AP: −2.0 mm, ML: −2.5 mm), and one occipital reference electrode (AP: −10.0 mm, ML: 0.0 mm). Additionally, two platinum electrodes were implanted to record LFP signals: one into the right mPFC (AP: +3.0 mm, ML: +0.5 mm, DV: −3.6 mm) and one into the right dHC (AP: −3.1 mm, ML: +3.0 mm, DV: −3.6 mm). Electrode positions were confirmed by histological analysis. One stainless steel wire electrode was implanted in the neck muscle for EMG recordings. Electrodes were connected to a six-channel electrode pedestal (PlasticsOne, USA) and fixed with cold polymerizing dental resin and the wound was sutured. Rats had at least 5 days for recovery.

**Recordings.** Rats were habituated to the recording box (dark gray PVC, 30 Å~ 30 cm, height: 40 cm) for two days, twelve hours per day. On the third day, animals were recorded for 12 hr, during the light phase, starting at 7:00 hr. The animal's behavior was continuously tracked using a video camera mounted on the recording box. EEG, LFP and EMG signals were continuously recorded and digitalized using a CED Power 1401 converter and Spike2 software (Cambridge Electronic Design). During the recordings, the electrodes were connected through a swiveling commutator to an amplifier (Model 15A54, Grass Technologies). The screw electrode in the occipital skull served as reference for all EEG, LFP, and EMG recordings. Filtering was for the EEG between 0.1 and 300 Hz, for LFP signals a high-pass filter of 0.1 Hz was applied, and for the EMG between 30 and 300 Hz. Signals were sampled at 1 kHz.

**Histology.** After the last recording session, electrolytic lesions were made at the tip of the electrodes to verify their precise location (dHC and mPFC). Rats were deeply anesthetized with a lethal dose of fentanyl, midazolam, and medetomidin and intracardially perfused with saline (0.9%, wt/vol) followed by a 4 per cent paraformaldehyde fixative solution. After extraction from the skull, brains were post-fixed in 4 per cent paraformaldehyde fixative solution for 1 day. Brains were then sliced into coronal sections (70 μm) and stained with 0.5 per cent toluidine blue.

**Sleep stage determination.** Sleep stages and wakefulness were determined offline based on EEG and EMG recordings, using standard visual scoring procedures for consecutive 10-s epochs as previously described (Neckelmann et al., 1994; Oyanedel et al., 2020; Table S4). Three sleep stages were discriminated: slow-wave sleep (SWS), intermediate stage (IS) and REM sleep. Wakefulness was identified by mixed-frequency EEG and sustained EMG activity, SWS by the presence of high amplitude low activity (delta activity: <4.0 Hz) and reduced EMG tone, REM sleep by low-amplitude EEG activity with predominant theta activity (5.0–10.0 Hz), phasic muscle twitches and decrease of EMG tone. IS was identified by a decreased delta activity, progressive increase of theta activity and presence of sleep spindles. Recordings were scored by two experienced experimenters (interrater agreement >89.9%). Afterward, consensus was achieved for epochs with divergent scoring.

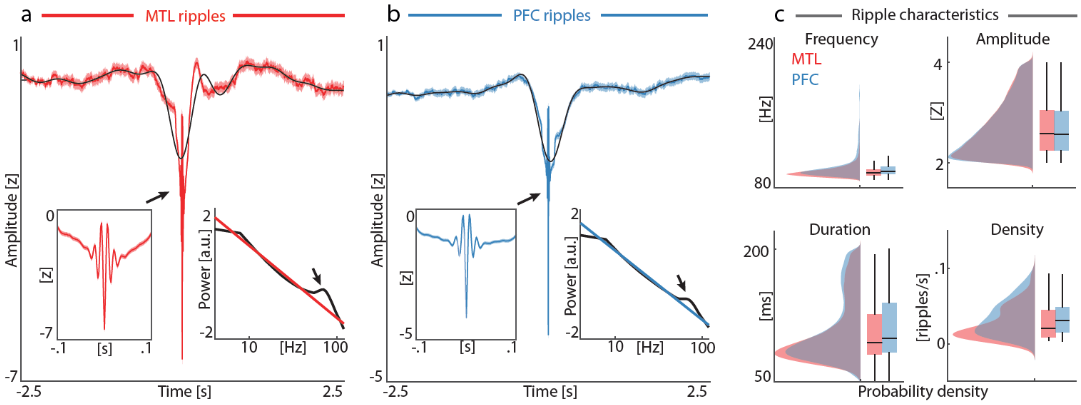


**Fig. S1. Ripple detection on broad frequency band (60-240 Hz). (a)** Group-level visualization of ripple events originating in the MTL (*N* = 45,891). Average ripple waveform (red trace) and superimposed low-pass filtered sharp wave (2 Hz; black trace). Insets highlight zoomed SW-R (*Left inset,* ± *.1s*) and power spectrum density relative to the ripple trough (±.1 s; black trace) including 1/f component (colored trace). **(b)** Ripple events originating in the PFC (*N* = 220,379) on group level using the same convention as in panel **a**. **(c)** Ripple characteristics for MTL and PFC channels. Note the peak frequencies for the MTL (90.25 ± .03, mean ± SEM) and PFC ripples (93.34 ± .02). Data are displayed as probability density functions; box plots represent median, $1^{st}/3^{rd}$ quartiles and extreme values

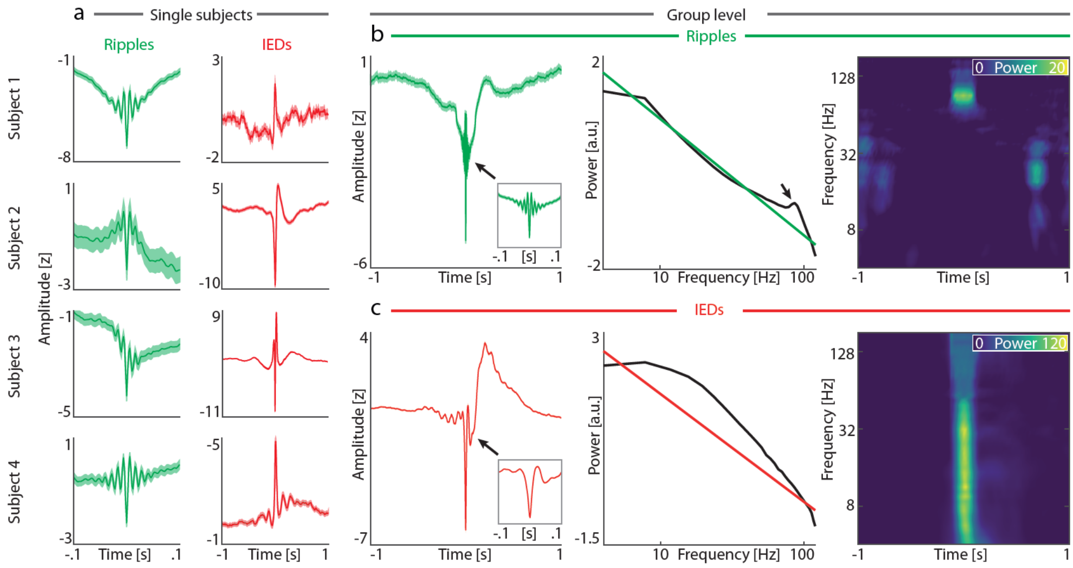


**Fig. S2.** Contrast between physiologic ripple events and pathologic interictal epileptiform discharges (IEDs) in the medial temporal lobe (MTL). **(a)** Single-subject examples of event-locked, broadband iEEG traces relative to ripples (*Left* column; green) and IEDs (*Right* column; red). **(b)** Group-level visualization of ripple events ($N_{channels}$ = 82; $N_{ripples}$ = 55,840). *Left*: Average ripple waveform. Inset shows the ripple oscillation for a shorter time window. *Middle*: Power spectrum density relative to the ripple trough (± .1 s; black) and aperiodic 1/f component (color). Note the peak in the power spectrum for the ripple frequency band. *Right*: Time-frequency spectrogram, showing a temporally-specific power increase for the ripple frequency band. **(c)** Group-level visualization of IEDs ($N_{channels}$ = 82; $N_{IEDs}$ = 116,456) using the same convention as in panel **b**. *Left*: Average IED waveform. Inset shows the IED waveform for a shorter time window. Note the predominant peak relative to ripple events. *Middle*: Power spectrum density relative to the IED peak (± .1 s; black trace) and aperiodic 1/f component (colored trace). Note the broadband increase in power. *Right*: Time-frequency spectrogram emphasizes the broadband increase in power upon IED occurrence.

**Table S1.** Subject descriptives

| Subject | Age | Gender | Handedness | English as first language | Recording Date | Recording Time | sEEG/Grid | MTL electrodes | PFC electrodes |
|---|---|---|---|---|---|---|---|---|---|
| 01 | 50 | F | **NA** | **NA** | 18/03/2016 | 21:28 - 08:28 | left sEEG, Grid | 7 | 37 |
| 02 | 42 | F | R | 0 | 26/07/2016 | 20:01 - 08:01 | sEEG, bilateral | 6 | 10 |
| 03 | 48 | F | R | 1 | 08/12/2016 | 22:28 - 07:02 | sEEG, bilateral | 4 | 15 |
| 04 | 24 | M | R | 1 | 13/12/2016 | 20:44 - 07:44 | sEEG, bilateral | 10 | 17 |
| 05 | 33 | M | R | **NA** | 05/01/2017 | 20:04 - 08:04 | sEEG, bilateral | 5 | 18 |
| 06 | 55 | F | L | 1 | 23/01/2017 | 19:47 - 08:30 | sEEG, bilateral | 4 | 15 |
| 07 | 33 | F | R | 1 | 22/02/2017 | 20:16 - 08:16 | sEEG | 5 | 26 |
| 08 | 23 | M | R | 1 | 22/03/2017 | 19:56 - 08:56 | sEEG, bilateral | 3 | 41 |
| 09 | 19 | F | R | 0 | 26/04/2017 | 20:22 - 07:43 | sEEG, bilateral | 2 | 12 |
| 10 | 50 | F | R | 0 | 20/09/2017 | 20:38 - 08:38 | sEEG, bilateral | 7 | 21 |
| 11 | 28 | M | R | 1 | 07/12/2017 | 21:55 - 07:55 | sEEG, bilateral | 4 | 20 |
| 12 | 58 | M | R | 1 | 24/01/2018 | 19:36 - 08:04 | sEEG, bilateral | 2 | 19 |
| 13 | 25 | F | R | 1 | 21/03/2018 | 19:41 - 07:41 | sEEG, bilateral | 8 | 49 |
| 14 | 27 | F | R | 1 | 19/06/2018 | 19:30 - 08:08 | sEEG, bilateral | 15 | 39 |

Notes: Age (years); Gender (F=Female; M=Male); Handedness (L=Left; R=Right); English as first Language (Y=Yes; N=No); Recording date (dd/mm/yyyy); Recording time (hh:mm); sEEG = stereo EEG; medial temporal lobe (MTL); prefrontal cortex (PFC); NA=Not Available.

**Table S2.** Sleep architecture – human data (mean ± SD)

| Subject | Sleep period [hh:mm] | Wake [%] | NREM1 [%] | NREM2 [%] | NREM3 [%] | REM [%] | Movement [%] |
|---|---|---|---|---|---|---|---|
| 01 | 08:41 | 19.94 | 7.57 | 50.43 | 7.67 | 12.66 | 1.73 |
| 02 | 10:11 | 18.23 | 6.87 | 24.04 | 36.14 | 13.16 | 1.55 |
| 03 | 08:37 | 43.04 | 2.61 | 37.04 | 10.25 | 6.96 | 0.1 |
| 04 | 11:42 | 16.8 | 7.05 | 46.55 | 12.6 | 16.87 | 0.14 |
| 05 | 12:45 | 44.77 | 6.73 | 34.18 | 2.48 | 11.5 | 0.33 |
| 06 | 10:05 | 26.42 | 7.93 | 47.98 | 12.8 | 4.71 | 0.17 |
| 07 | 12:20 | 49.32 | 4.32 | 27.09 | 5.61 | 13.65 | 0 |
| 08 | 13:49 | 33.94 | 7.96 | 29.48 | 12.6 | 15.55 | 0.48 |
| 09 | 13:49 | 33.94 | 7.96 | 29.48 | 12.6 | 15.55 | 0.48 |
| 10 | 12:16 | 11.81 | 3.8 | 48.13 | 28.04 | 8.15 | 0.07 |
| 11 | 10:12 | 35.54 | 16.75 | 29.98 | 11.44 | 0 | 6.29 |
| 12 | 07:22 | 58.82 | 8.82 | 26.58 | 2.04 | 3.39 | 0.34 |
| 13 | 10:55 | 30.21 | 13.73 | 36.99 | 14.11 | 3.66 | 1.3 |
| 14 | 09:50 | 10.93 | 4.24 | 58.56 | 14.15 | 11.36 | 0.76 |
| **Group average:** | 10:54 ± 01:58 | 30.98 ± 14.05 | 7.60 ± 3.65 | 37.61 ± 10.42 | 13.04 ± 8.82 | 9.80 ± 5.12 | 0.98 ± 1.57 |

**Table S3.** Ripple descriptives – human data; group level (mean ± SD)

| Region | Subjects | Electrodes | State | Ripples [#] | Frequency [Hz] | Amplitude [z] | Duration [ms] | Sharp transient [z] | Density [events/s] |
|---|---|---|---|---|---|---|---|---|---|
| MTL | 14 | 86 | Wake | 42144 | 90.962 ± 7.356 | 2.681 ± 0.511 | 108.555 ± 35.095 | 1.481 ± 0.334 | 0.033 ± 0.024 |
| | | | NREM1 | 7820 | 90.366 ± 6.973 | 2.692 ± 0.517 | 109.023 ± 35.229 | 1.453 ± 0.339 | 0.033 ± 0.024 |
| | | | NREM2 | 43514 | 90.578 ± 7.001 | 2.685 ± 0.514 | 108.888 ± 35.186 | 1.426 ± 0.349 | 0.032 ± 0.023 |
| | | | NREM3 | 12265 | 90.953 ± 7.435 | 2.685 ± 0.516 | 107.503 ± 34.262 | 1.390 ± 0.357 | 0.029 ± 0.023 |
| | | | REM | 9671 | 90.942 ± 7.678 | 2.630 ± 0.491 | 107.954 ± 34.606 | 1.445 ± 0.346 | 0.031 ± 0.022 |
| PFC | 14 | 368 | Wake | 228188 | 94.136 ± 11.458 | 2.682 ± 0.504 | 109.901 ± 35.783 | 1.532 ± 0.318 | 0.040 ± 0.027 |
| | | | NREM1 | 46334 | 93.898 ± 11.172 | 2.678 ± 0.506 | 110.771 ± 36.027 | 1.516 ± 0.323 | 0.040 ± 0.025 |
| | | | NREM2 | 197384 | 93.162 ± 11.018 | 2.689 ± 0.513 | 110.436 ± 36.166 | 1.479 ± 0.341 | 0.040 ± 0.025 |
| | | | NREM3 | 61942 | 93.584 ± 11.207 | 2.686 ± 0.515 | 111.045 ± 36.193 | 1.412 ± 0.364 | 0.037 ± 0.024 |
| | | | REM | 47763 | 92.902 ± 11.253 | 2.699 ± 0.513 | 109.168 ± 35.620 | 1.517 ± 0.324 | 0.038 ± 0.026 |

Notes: Medial temporal lobe (MTL); Prefrontal cortex (PFC).

**Table S4.** Sleep architecture – rodent data (mean ± SD)

| Rodent | Sleep period [hh:mm] | Wake [%] | SWS [%] | Intermediate stage (IS) [%] | REM [%] |
|---|---|---|---|---|---|
| 01 | 11:27 | 36.83 | 51.55 | 1.43 | 10.16 |
| 02 | 11:29 | 37.08 | 48.27 | 3.36 | 11.24 |
| 03 | 11:41 | 28.14 | 57.28 | 1.83 | 12.75 |
| 04 | 11:07 | 28.68 | 58.24 | 2.2 | 10.86 |
| Group average: | 11:26 ± 00:14 | 32.68 ± 4.28 | 53.84 ± 4.11 | 2.21 ± 0.72 | 11.25 ± 0.95 |

**Table S5.** Ripple descriptives – rodent data; group level (mean ± SD)

| ROI | Rodents | Electrodes | State | Ripples [#] | Frequency [Hz] | Amplitude [z] | Duration [ms] | Sharp transient [z] | Density [events/s] |
|---|---|---|---|---|---|---|---|---|---|
| MTL | 4 | 4 | Wake | 1091 | 105.371 ± 14.282 | 2.665 ± 0.509 | 110.947 ± 34.312 | 1.410 ± 0.352 | 0.017 ± 0.013 |
| | | | SWS | 3037 | 99.335 ± 11.582 | 2.708 ± 0.522 | 114.749 ± 35.460 | 1.438 ± 0.332 | 0.035 ± 0.028 |
| | | | REM | 624 | 94.343 ± 8.656 | 2.579 ± 0.468 | 103.128 ± 33.310 | 1.436 ± 0.314 | 0.035 ± 0.025 |
| | | | pre-REM | 92 | 94.348 ± 9.868 | 2.604 ± 0.471 | 106.791 ± 36.886 | 1.472 ± 0.325 | 0.028 ± 0.023 |
| PFC | 4 | 4 | Wake | 1276 | 111.011 ± 18.531 | 2.847 ± 0.579 | 104.642 ± 32.674 | 1.267 ± 0.377 | 0.020 ± 0.009 |
| | | | SWS | 877 | 98.603 ± 12.001 | 2.514 ± 0.460 | 104.129 ± 30.871 | 1.342 ± 0.365 | 0.010 ± 0.009 |
| | | | REM | 191 | 102.251 ± 11.569 | 2.328 ± 0.368 | 107.111 ± 32.114 | 1.428 ± 0.327 | 0.011 ± 0.014 |
| | | | pre-REM | 15 | 100.667 ± 11.782 | 2.483 ± 0.445 | 100.350 ± 22.910 | 1.455 ± 0.359 | 0.004 ± 0.005 |
